# A comparison of non-magnetic and magnetic beads for measuring IgG antibodies against *P. vivax* antigens in a multiplexed bead-based assay using Luminex^®^ technology (Bio-Plex^®^ 200 or MAGPIX^®^)

**DOI:** 10.1101/2020.08.10.243980

**Authors:** Ramin Mazhari, Jessica Brewster, Rich Fong, Caitlin Bourke, Zoe SJ Liu, Eizo Takashima, Takafumi Tsuboi, Wai-Hong Tham, Matthias Harbers, Chetan Chitnis, Julie Healer, Maria Ome-Kaius, Jetsumon Sattabongkot, James Kazura, Leanne J. Robinson, Christopher King, Ivo Mueller, Rhea J. Longley

**Author notes:** Corresponding author (RJL). These authors contributed equally to this work (Ivo Mueller, Rhea J. Longley).

## Abstract

Multiplexed bead-based assays that use Luminex xMAP^®^ technology have become popular for measuring antibodies against proteins of interest in many fields, including malaria and more recently SARS-CoV-2/COVID-19. There are currently two formats that are widely used: non-magnetic beads or magnetic beads. Data is lacking regarding the comparability of results obtained using these two types of beads, and for assays run on different instruments. Whilst non-magnetic beads can only be run on flow-based instruments (such as the Luminex^®^ 100/200™ or Bio-Plex^®^ 200), magnetic beads can be run on both these and the newer MAGPIX^®^ instruments. In this study we utilized a panel of purified recombinant *Plasmodium vivax* proteins and samples from malaria-endemic areas to measure *P. vivax*-specific IgG responses using different combinations of beads and instruments. We directly compared: i) non-magnetic versus magnetic beads run on a Bio-Plex^®^ 200, ii) magnetic beads run on the Bio-Plex^®^ 200 versus MAGPIX^®^ and iii) non-magnetic beads run on a Bio-Plex^®^ 200 versus magnetic beads run on the MAGPIX^®^. We also performed an external validation of our optimized assay. We observed that IgG antibody responses, measured against our panel of *P. vivax* proteins, were strongly correlated in all three of our comparisons, however higher amounts of protein were required for coupling to magnetic beads. Our external validation indicated that results generated in different laboratories using the same coupled beads are also highly comparable, particularly if a reference standard curve is used.

## Introduction

Over the past 5-10 years there has been a rapid uptake of Luminex bead-based technologies to measure antibody responses to multiple proteins simultaneously. These assays have numerous advantages over traditional enzyme-linked immuosorbent assays (ELISA), such as a reduction in sample volume required and reduced laboratory time, as well as the main advantage of allowing multiplexed detection of antibody responses. This is particularly relevant for the detection of antibodies against complex pathogens that express many hundreds to thousands of proteins, such as the *Plasmodium* parasites (the causative agent of malaria). Access to standardized control reagents [1] will also allow results from these assays to be reliably compared between different laboratories, which may result in more consistent findings between different studies [2].

Multiplexed bead-based assays use Luminex xMAP^®^ technology [3], which centers on use of beads (microspheres) with different fluorescent colours that can be detected in unique regions on a compatible instrument such as a Luminex^®^ 200™ (also known as a Bio-Plex^®^ 200, sold by Bio-Rad) or MAGPIX^®^. Proteins of interest can be coupled to a unique set of beads, facilitating multiplexed detection of antibody responses to multiple proteins. Several studies have been conducted with a focus on optimizing various steps of the coupling process or assay work-flow, in the context of detection of antibodies against *Plasmodium* proteins, such as bead coupling [4], sample pre-dilution [4], assay temperature [4], plate washing [4], operator expertise [4], incubation times [1], and bead numbers [5]. Two different types of beads are available for coupling proteins: non-magnetic and magnetic. Non-magnetic beads can only be run on flow-based instruments such as the Luminex^®^ 200™/Bio-Plex^®^ 200, whilst magnetic beads can be run on both, flow-based instruments and the MAGPIX^®^. The MAGPIX^®^ is based on CCD imaging technology, and offers advantages over the flow-based systems such as faster acquisition time, reduced use of reagents such as sheath fluid and the reduced cost of the MAGPIX^®^ instrument compared to the Luminex^®^ 200™/Bio-Plex^®^ 200 instruments.

The primary aim of this study was to perform a series of comparisons of both non-magnetic and magnetic beads and assaying those beads on the Bio-Plex^®^ 200 or the MAGPIX^®^. A secondary aim was to demonstrate that this assay is highly reproducible in an independent laboratory through an external validation. This study used a panel of 19 different *P. vivax* proteins and plasma samples from *P. vivax*-endemic areas to detect *P. vivax*-specific IgG responses, however the large number of proteins assessed and consistent results obtained, suggest these findings should be generalizable for optimization of the multiplexed bead-based assay for other pathogens. This is important in the context of the ongoing SARS-CoV-2 pandemic, as multiple laboratory assays based on Luminex technology are under development [6–8].

## Materials and methods

### Plasma samples

For all assays described here, a pool of samples from individuals from Papua New Guinea (PNG) with high levels of anti-*Plasmodium* antibodies was used as a positive control for the standard curve dilution to adjust for plate to plate variation, as previously described [9].

Two sets of plasma samples from malaria-endemic areas were used for comparisons of non-magnetic and magnetic beads, and the different acquisition instruments. These were 80 individuals from a longitudinal observational cohort study in Thailand, conducted in the Kanchanaburi and Ratchaburi provinces in 2013-2014. This cohort has previously been described in detail [10, 11], and the 80 plasma samples used were collected at the last visit of the cohort. The second set of samples came from a longitudinal observational cohort study in the Solomon Islands, conducted on the island Ngella in 2013-2014. This cohort has previously been described in detail [10, 12], and 83 plasma samples were used from individuals at the last visit of this cohort.

An additional set of plasma samples from a cohort study in PNG was used for external validation of the assay. Samples were selected from the Mugil II paediatric cohort study. The study enrolled 450 children aged 5-12 years old in 2012 from the Mugil area on the North Coast of Madang province. All children were given antimalarial drugs to eliminate blood-stage *Plasmodium spp* and blood samples were collected for parasitological and immunological studies. For the external validation, a set of 425 samples was used from the baseline timepoint (collected 2 weeks after drug treatment).

### Ethics statement

All samples were collected after approval from local ethics committees, with volunteers/participants providing written informed consent and/or assent. The Ethics Committee of the Faculty of Tropical Medicine, Mahidol University, Thailand approved the Thai cohort study (MUTM 2013-027-01). The National Health Research and Ethics Committee of the Solomon Islands Ministry of Health and Medical Services (HRC12/022) approved the Solomon Islands study. The Mugil II paediatric cohort was approved by the PNG Institute of Medical Research Institutional Review Board (IMR IRB) (1116/1204), the PNG Medical Research Advisory Committee (MRAC) (11.21/1206), the Walter and Eliza Hall Institute Human Research Ethics Committee (WEHI HREC) (12/09), and the Case Western Reserve University Hospitals of Cleveland Medical Center (CWRU UHCMC) (05-11-11). The HREC at WEHI approved samples for use in Melbourne (#14/02).

### Coupling *P. vivax* proteins to non-magnetic and magnetic beads

The carboxylated beads were sourced from Bio-Rad (Bio-Plex COOH Beads, 1ml, 1.25×10^7^ beads/ml and Bio-Plex Pro Magnetic COOH Beads, 1ml, 1.25×10^7^ bead/ml) and stored at 2-4°C. Optimisation of coupling procedures for non-magnetic and magnetic beads were done separately, due to the larger size of the magnetic beads generally requiring more protein (see Results). To be able to measure all plasma samples at the same dilution, we optimized all protein concentrations by generating a log-linear standard curve with a positive control plasma pool from immune PNG donors (high responders to *Plasmodium* antigens).

Coupling of *P. vivax* proteins to non-magnetic beads was performed as previously described [10]. Briefly, the optimised antigen concentration (Tables 1 and 2) was coupled to 2.5×10^6^ pre-activated microspheres, in 100 mM monobasic sodium phosphate buffer pH 6.0, using 50mg/ml sulfo-NHS and 50 mg/ml of EDC to cross-link the proteins to the beads. The activated beads were washed and stored in PBS, 0.1% BSA, 0.02% Tween-20, 0.05% Na-azide, pH 7.4 at 4°C until use. For the coupling to magnetic beads, a magnet rack was used for pelleting the beads, instead of the centrifugation step for non-magnetic beads. We qualitatively assessed the stability of the coupled beads by visual comparison of the MFI of the standard curve over a nine-month period.

**Table 1:** *P. vivax* proteins used in the comparison experiments, with the amount of protein coupled per non-magnetic and magnetic beads indicated. Gene annotations and protein IDs were sourced from PlasmoDB (release 36, http://plasmodb.org/plasmo/), or GenBank when necessary.

| Gene Annotation | Protein ID | Expression System | Protein Concentration (µg/ul) | Construct, amino acids (size) | Protein amount (µg/1x10<sup>6</sup>) non-magnetic beads | Protein amount (µg/1x10<sup>6</sup>) magnetic beads |
| --- | --- | --- | --- | --- | --- | --- |
| RBP2b (P25) | PVX_094255 | *E. coli* | 4.15 | 161-1454 (1294) | 0.21 | 0.24 |
| MSP1-19 | PVX_099980 | WGCF | 1.55 | 1622-1729 (108) | 0.30 | 1.60 |
| RBP2b | PVX_094255 | WGCF | 2.06 | 1986-2653 (667) | 0.28 | 3.20 |
| RAMA | PVX_087885 | WGCF | 0.78 | 462-730 (269) | 0.06 | 0.48 |
| PvEBPII | KMZ83376.1 | *E. coli* | 10 | 109-432 (324) | 0.08 | 0.20 |
| SSA-s16 | PVX_000930 | WGCF | 0.41 | 31-end (110) | 0.40 | 0.80 |
| PvRIPR | PVX_095055 | *E. coli* | 1 | 552-1075 (524) | 0.40 | 0.80 |
| MSP3.10 | PVX_097720 | WGCF | 0.64 | 25-end (828) | 0.40 | 0.80 |
| Hyp. Protein | PVX_097715 | WGCF | 0.7 | 20-end (431) | 0.14 | 1.20 |
| PvDBPII (AH) | AAY34130.1 | *E. coli* | 0.6 | 1-237 (237) | 0.43 | 0.56 |
| MSP8 | PVX_097625 | WGCF | 0.39 | 24-463 (440) | 0.28 | 0.56 |
| Unspecified/<br>Pv-fam-a | PVX_112670 | WGCF | 1.13 | 34-end (302) | 0.45 | 0.90 |
| Pv-fam-a | PVX_096995 | WGCF | 1.7 | 61-end (420) | 0.34 | 1.20 |
| MSP3.3 | PVX_097680 | WGCF | 0.55 | 21-end (996) | 0.48 | 0.32 |
| MSP7.1 | PVX_082700 | WGCF | 0.33 | 23-end (397) | 0.40 | 0.60 |
| MSP5 | PVX_003770 | WGCF | 0.58 | 23-365 (343) | 0.01 | 0.016 |
| MSP7 | PVX_082670 | WGCF | 0.61 | 24-end (388) | 0.40 | 0.40 |
| PvTRAP/<br>SSP2 | PVX_082735 | WGCF | 0.9 | 26-493 (468) | 0.40 | 0.80 |
| PvDBPII (sal1) | PVX_110810 | *E. coli* | 1.2 | 193-521 (329) | 0.29 | 0.24 |

**Table 2:** *P. vivax* proteins used for the external validation. Proteins were coupled to non-magnetic beads at WEHI and half of each batch of bead-conjugated protein was shipped to CWRU.

| Gene Annotation | Protein ID | Expression System |
| --- | --- | --- |
| MSP1-19 | PVX_099980 | WGCF |
| Pv-fam-a | PVX_096995 | WGCF |
| hypothetical protein, conserved | PVX_094830 | WGCF |
| Pv-fam-a | PVX_112670 | WGCF |
| MSP7 | PVX_082650 | WGCF |
| RBP2b | PVX_094255 | WGCF |
| hypothetical protein, conserved | PVX_001000 | WGCF |
| merozoite surface protein 8 | PVX_097625 | WGCF |
| PvTRAP/SSP2 | PVX_082735 | WGCF |
| MSP7 | PVX_082645 | WGCF |
| PvRBP-2, putative | PVX_090330 | WGCF |
| sexual stage antigen s16,<br>putative | PVX_000930 | WGCF |

*Plasmodium vivax* recombinant antigens were expressed and purified in three countries: Japan (Takafumi Tsuboi, Ehime University & Matthias Harbers, CellFree Sciences), Australia (Wai-Hong Tham and Julie Healer, Walter & Eliza Hall Institute of Medical Research) and France (Chetan Chitnis, Institut Pasteur). Proteins were expressed either in the wheat-germ cell-free expression system (WGCF) or *E. coli.* See Table 1 for a complete list of proteins and the optimised amount coupled to non-magnetic and magnetic beads.

### Multiplexed assay for measurement of *P. vivax-*specific antibody responses

To measure the IgG levels, a multiplexed bead based assay was used, as previously described [10]. Briefly, antigen-specific IgG was detected by incubating 500 beads of each antigen per well with plasma diluted at 1:100, in a final volume of 100μl. Non-magnetic beads were washed using a vacuum manifold, whereas magnetic beads were washed using a magnetic plate washer. After the washings, a 1:100 dilution of PE-conjugated Donkey F(ab)2 anti-human IgG (JIR 709-116-098) was added. At least 15 beads of each region/antigen were then acquired and analysed on a Bio-Plex^®^ 200 instrument and/or a MAGPIX^®^ instrument as per the manufacturer’s instructions. Note that for comparing data between Bio-Plex^®^ 200 and MAGPIX^®^ instruments it is important that the “high RP1” target is not selected on the Bio-Plex^®^ 200, as this option is not available on the MAGPIX^®^. On each plate, a twofold serial dilution from 1/50 to 1/25,600 of a seropositive control plasma pool (generated from PNG adults) was included. Note that for the external validation both labs used the same PNG control pool to generate the standard curve.

The results were expressed as mean fluorescence intensity (MFI) of at least 15 beads for each antigen.

### Instruments

Antibody measurements were acquired using a Bio-Plex^®^ 200 Multiplexing Analyzer System from Bio-Rad for all non-magnetic coupled beads (Bio-Plex^®^ 200System, Bio-Plex^®^ high-throughput fluidics system, microplate platform and a computer with the Bio-Plex® manager software v.5.0). Washing steps were carried out on a Bio-Rad Aurum vacuum manifold.

For all magnetic coupled beads a MAGPIX^®^ Multiplexing System from Millipore was used (MAGPIX^®^ System and the Xponent software V.4.2). Washing steps were carried out using a magnetic plate washer from BioTek Instruments (BioTek ELx50). A Bio-Rad Sure Beads magnetic rack was used during the coupling process.

Plates were incubated on a Ratek Platform shaker (Microtiter/PCR Plate Shaker). A Vortex Sonicator (Branson 2200), a BioSan Vortex V-1 plus and a Table centrifuge (Eppendorf Centrifuge 5424) were also used during the coupling process.

### Statistical analysis

The raw MFI results were converted to relative antibody units (RAU) using protein-specific standard curve data. A log–log model was used to obtain a more linear relationship, and a five-parameter logistic function was used to obtain an equivalent dilution value compared to the PNG control plasma (ranging from 1.95×10^−5^ to 0.02). The interpolation was performed in R. Pearson’s r^2^ correlations were performed to determine the strength of correlation and the statistical significance for all comparisons. To enable these parametric correlations, data were log-transformed prior to the analysis to better fit the normal distribution.

## Results and discussion

### Comparison of total IgG antibodies detected against *P. vivax* antigens coupled to either non-magnetic or magnetic beads and assayed by a Bio-Plex^®^ 200 instrument

Total IgG antibody levels against a panel of 19 *P. vivax* proteins, measured in plasma samples from 163 individuals living in malaria-endemic areas of Thailand and the Solomon Islands, were assayed using either non-magnetic or magnetic beads and run on a Bio-Plex^®^ 200 instrument. IgG levels to 18 of 19 proteins were well correlated between non-magnetic and magnetic assays, with Pearson r^2^-values ranging from 0.29-0.95 (all p<0.0001) (Figure 1), supporting previous findings based on *P. falciparum* proteins [13]. This is despite different amounts of each protein being coupled to non-magnetic versus magnetic beads (Table 1). The exception was for the protein PVX_003770 (MSP5), with the lowest correlation coefficient at r^2^=0.075 (p<0.001). A sub-set of the samples that had relatively high antibody levels for PVX_003770 when assayed with non-magnetic beads had relatively low antibody levels when assayed with magnetic beads, likely accounting for the lower correlation coefficient observed. Interestingly, the amount of protein coupled for PVX_003770 (for both non-magnetic and magnetic beads) was substantially lower than for the other proteins. Future experiments are planned to determine whether increasing the protein amount for PVX_003770 could result in a higher correlation between the two platforms.

**Figure 1:**
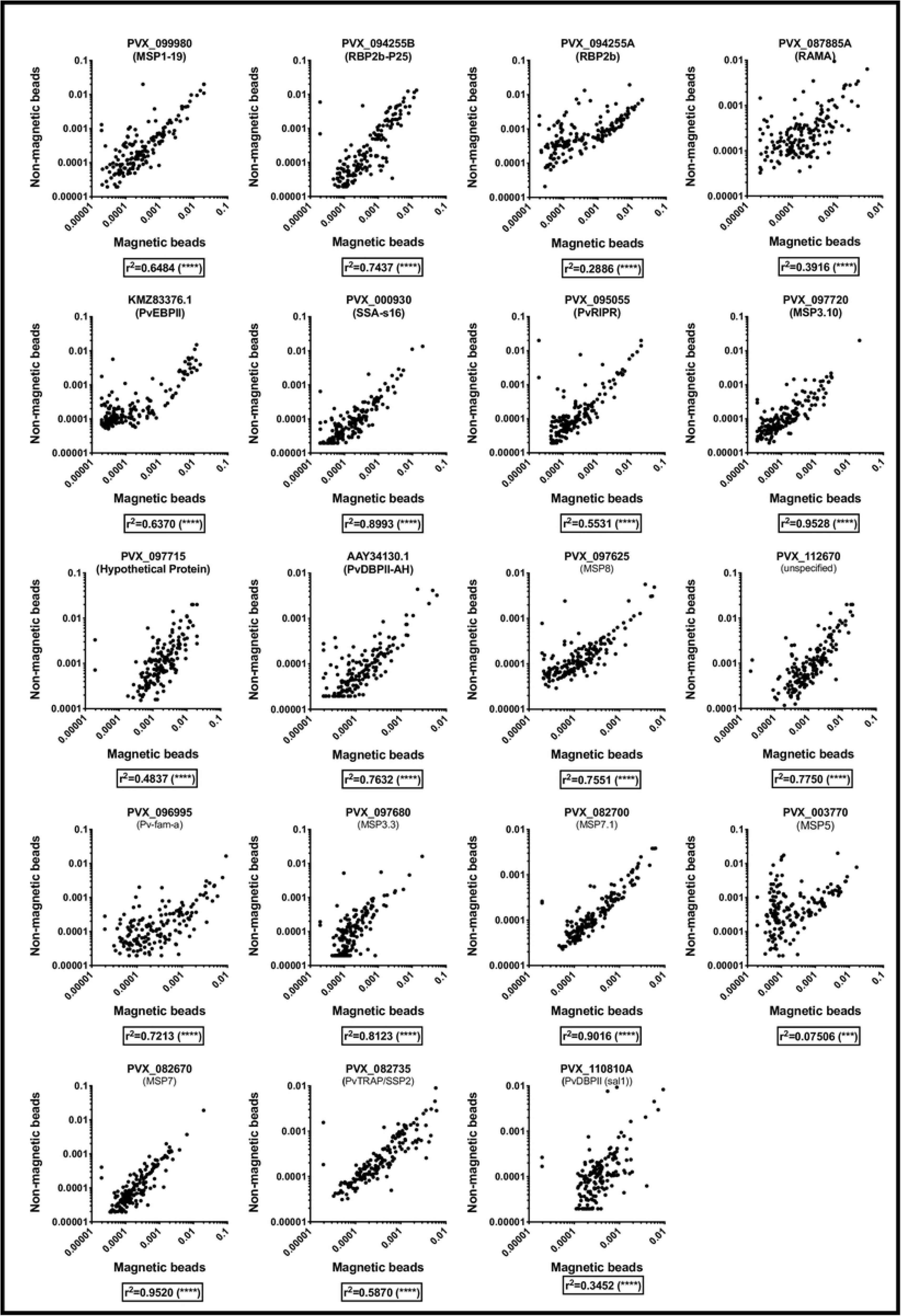
IgG antibody levels (RAU) measured against 19 *P. vivax* proteins in samples from malaria-endemic areas, using either non-magnetic or magnetic beads and run on a Bio-Plex^®^ 200 instrument. *** p<0.001, **** p<0.0001.

### Comparison of total IgG antibodies detected against *P. vivax* antigens coupled to magnetic beads and assayed using either a Bio-Plex^®^ 200 instrument or a MAGPIX^®^ instrument

For this comparison, all 19 *P. vivax* antigens were coupled to magnetic beads only, at the optimised antigen concentrations. Total IgG antibody levels were measured in the same set of 163 plasma samples, with the assay run on both a Bio-Plex^®^ 200 and a MAGPIX^®^ instrument. To our knowledge, this is the first published report of this comparison. Here, the Pearson r^2^ correlation coefficients indicated a high level of correlation between samples run on both instruments (r^2^=0.970-0.999, p<0.0001, Figure 2). These results indicate that results obtained on either platform, when antigens are coupled at the same optimised concentrations to magnetic beads, are highly comparable. The strength of the correlations in this comparison is stronger than the previous analysis (which compared non-magnetic versus magnetic beads on the same instrument), presumably because the same sets of coupled beads were run on both instruments. The strength of the correlations suggests that results obtained on the Bio-Plex^®^ 200 and MAGPIX^®^ are interchangeable.

**Figure 2:**
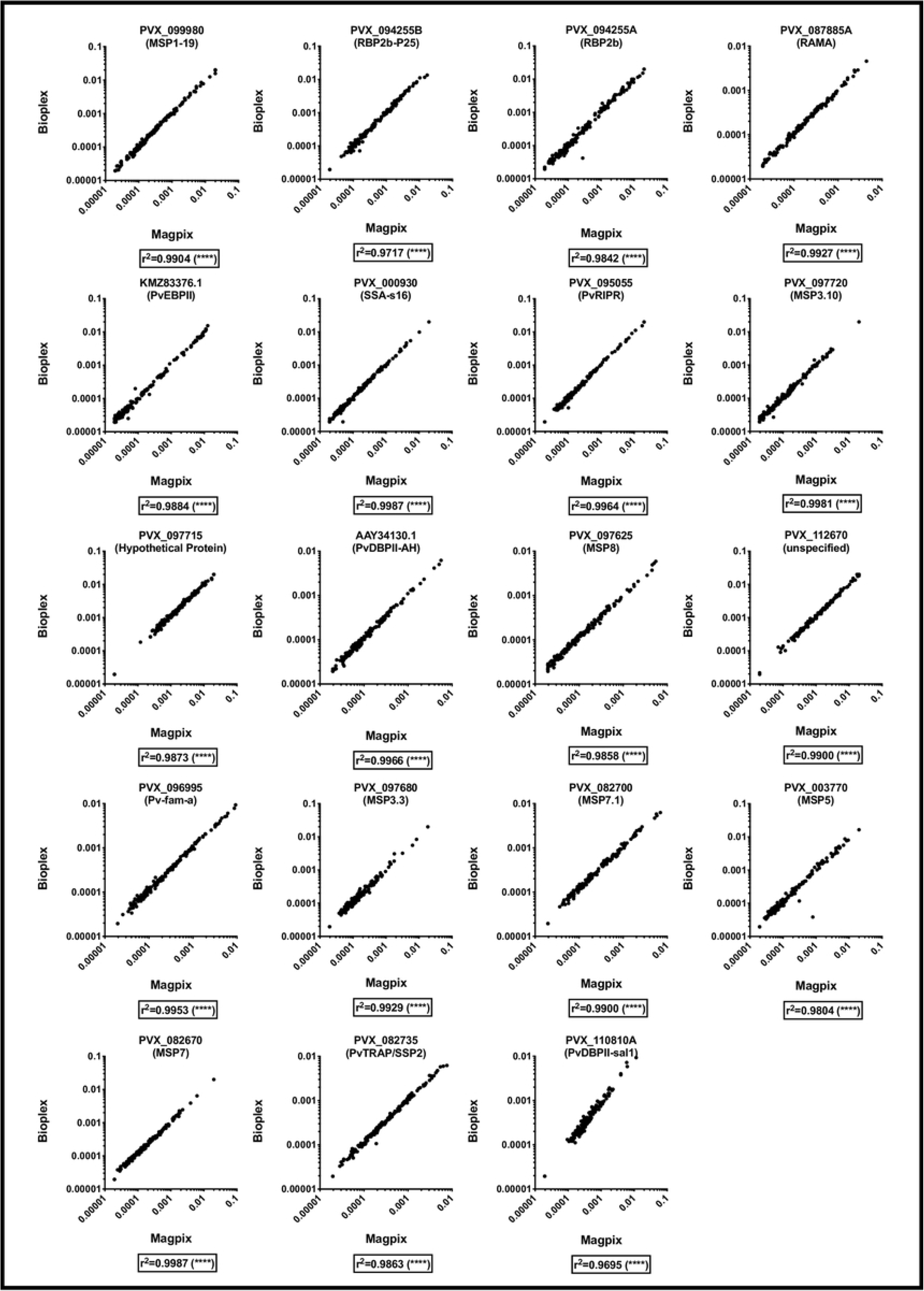
IgG antibody levels (RAU) measured against 19 *P. vivax* proteins in samples from malaria-endemic areas, using magnetic beads and run on either a Bio-Plex^®^ 200 instrument or MAGPIX^®^ instrument. **** p<0.0001.

### Comparison of total IgG antibodies against *P. vivax* antigens coupled to non-magnetic beads and analyzed on a Bio-Plex^®^ 200 instrument and antigens coupled to magnetic beads and analyzed on a MAGPIX^®^ instrument

The final comparison we wanted to conduct was of antigens coupled to non-magnetic beads and assayed on a Bio-Plex^®^ 200 instrument with antigens coupled to magnetic beads and assayed on a MAGPIX^®^ instrument. As non-magnetic beads are cheaper to purchase, users that have only a Bio-Plex^®^ 200 instrument would potentially favour this configuration (even though the instrument can run both non-magnetic and magnetic beads). Conversely, for users that only have a MAGPIX^®^ instrument, they are only able to run magnetic beads as the instrument cannot detect non-magnetic beads. To our knowledge, this is the first published report of this comparison for a non-commercial assay.

It was again observed that there was a strong correlation between results obtained using the non-magnetic beads/Bio-Plex^®^ 200 and magnetic beads/MAGPIX^®^ platforms, with Pearson r^2^ correlation coefficients ranging from 0.18-96 (p<0.0001, Figure 3). These correlation coefficients are similar to those obtained in the first comparison (non-magnetic versus magnetic beads both run on the Bio-Plex^®^ 200 instrument), and provide further support for our finding that antigens coupled to either type of beads and run on either instrument generally give very comparable total IgG measurements. As we observed in the first comparison, the weakest correlation was again for the protein PVX_003770 (r^2^=0.18).

**Figure 3.**
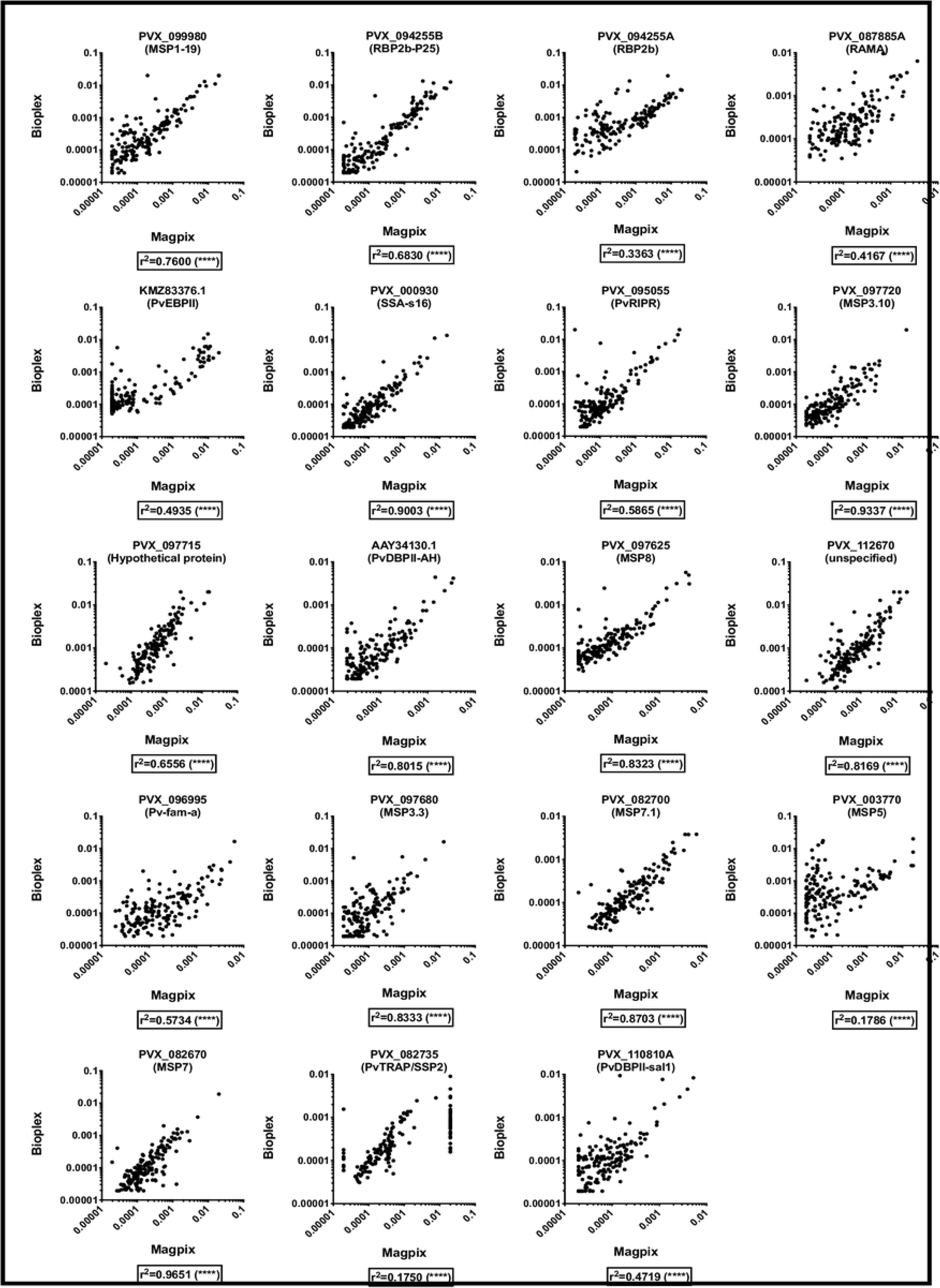
IgG antibody levels (RAU) measured against 19 *P. vivax* proteins in samples from malaria-endemic areas, using non-magnetic beads and run on a Bio-Plex^®^ 200 compared to use of magnetic beads run on a MAGPIX^®^ instrument. **** p<0.0001.

### External validation of a multiplexed assay using *P. vivax* antigens coupled to non-magnetic beads and analyzed on a Bio-Plex^®^ 200 instrument

The results thus far indicate that IgG levels measured using either non-magnetic or magnetic beads and assayed on either a Bio-Plex^®^ 200 or MAGPIX^®^ instrument are highly comparable. A group of 3 staff members, but all at the same Institute (Walter & Eliza Hall Institute, WEHI) using the same instruments, performed these measurements. Therefore an additional comparison was performed: external validation of the assay at an independent research Institute located overseas (Case Western Reserve University, CWRU).

A set of 425 plasma samples were aliquoted at CWRU and shared with WEHI. At the same time, a set of 12 *P. vivax* proteins (Table 2) were coupled to non-magnetic beads at WEHI and shared with CWRU. During the same week assays were performed to measure total IgG antibodies against these *P. vivax* antigens in the 425 plasma samples on Bio-Plex^®^ 200 instruments independently at each Institute (total of 6 plates run at each Institute). After exclusion of plates or samples following quality control checks (positive control – log-linear standard curve; bead counts > 15), data from 318 samples was directly compared between sites. The drop from 425 to 318 samples was largely due to one plate with failed standard curves that could not be repeated due to sample availability. IgG levels were compared first using raw data (MFI values). The Pearson r^2^ correlation coefficients indicated a strong correlation for all proteins with r^2^-values > 0.58 (p<0.0001), with the exception of PVX_094255 (RBP2b) (r^2^=0.32, p<0.0001) (Table 3, scatter plots in Figure S1). The same correlation analysis was then performed on data converted in R using the standard curves (to account for any plate-plate variation). Strong correlation coefficients were observed for all 12 proteins, including PVX_094255 (r^2^ values >0.51, p<0.0001) (Table 3, scatter plots in Figure S2). For the majority of proteins, the correlation was stronger after conversion (Table 3). This is expected given the conversion, based on the standard curve generated with a plasma pool from immune PNG donors, is used to account for any plate-plate variation.

**Table 3:** External validation of the non-magnetic bead assay run on the Bio-Plex^®^ 200. Pearson r^2^ correlation coefficients are shown for both the raw data (MFI) and the standard curve converted data (RAU). **** p<0.0001.

| Protein ID | Correlation MFI (n=318) | Correlation RAU (n=318) |
| --- | --- | --- |
| PVX_099980 | 0.76 **** | 0.85 **** |
| PVX_096995 | 0.69 **** | 0.76 **** |
| PVX_094830 | 0.58 **** | 0.55 **** |
| PVX_112670 | 0.63 **** | 0.65 **** |
| PVX_082650 | 0.73 **** | 0.70 **** |
| PVX_094255 | 0.32 **** | 0.51 **** |
| PVX_001000 | 0.69 **** | 0.70 **** |
| PVX_097625 | 0.73 **** | 0.73 **** |
| PVX_082735 | 0.81 **** | 0.85 **** |
| PVX_082645 | 0.79 **** | 0.75 **** |
| PVX_090330 | 0.71 **** | 0.66 **** |
| PVX_000930 | 0.80 **** | 0.83 **** |

These results indicate that data generated using this multiplexed assay are highly reproducible in a different laboratory setting when the same coupled-beads are used, particularly if both laboratories have access to the same positive control for standardization. Unfortunately, whilst there is a WHO reference reagent for *P. falciparum* serology studies [14], there is not yet a similar product available for *P. vivax.* Importantly, we also assessed the stability of the coupled beads by running the standard curve 10 times over a period of 9 months (intensely for 2 months) (Figure S3). For most proteins the coupled beads were highly stable (11/16 tested over 9-months), with the MFI dropping for three proteins and increasing for two proteins. This is supported by previous research that has indicated the stability of protein-coupled beads [13], noting that the stability may vary by antigen [15].

## Conclusions

The aim of this study was to demonstrate that multiplexing assays performed using magnetic beads or non-magnetic beads are highly comparable, independent of the beads and platform used to analyze the assays. We compared here a total of 19 *P. vivax* proteins that were coupled to both magnetic beads and non-magnetic beads. The protein concentration used for the couplings was individually determined by optimisation for each protein for the chosen bead type (Table 1). For this, a dilution series from the positive control plasma pool, prepared from immune PNG donors, was used to generate a log-linear standard curve for each protein. The non-magnetic beads are 5.5μm in size, whilst the magnetic beads are 6.5μm in size, likely accounting for the need to couple on average 0.3μg of protein to non-magnetic versus 0.8 μg of protein to magnetic beads (per 1×10^6^ beads). One coupling reaction using these amounts of protein is enough to assay > 3000 samples in singlicate, thus the slightly higher amount of protein required for magnetic beads is unlikely to be a limitation to using this format. We did not assess the efficiency of antigen coupling, which could potentially be an important variable impacting the amount of protein required for coupling.

We have demonstrated that results are highly comparable whether using proteins coupled to magnetic beads or non-magnetic beads and analysed using either a Bio-Plex^®^ 200 (non-magnetic and magnetic beads) or MAGPIX^®^ (magnetic beads only). Our external validation has also demonstrated that results generated in different laboratories are highly comparable, if a reference standard curve is included for standardization. Therefore researchers can, in principle, compare data generated with a different type of bead or assayed using a different instrument platform, if the amount of protein coupled is optimised for the correct type of bead. Overall, the choice of assay platform and instrument used is up to the user. Table 4 lists a number of factors that differ between the two platforms that users should consider. An important consideration is that up to 100 different proteins can be assayed simultaneously using non-magnetic beads and a Bio-Plex^®^ 200 instrument, whereas the maximum is 50 proteins using a MAGPIX^®^. If less than 50 proteins will be used, the MAGPIX^®^ instrument is cheaper and enables washing steps to be conducted with magnets, which improves both bead retention [13, 16] and speed of the assay.

**Table 4:**
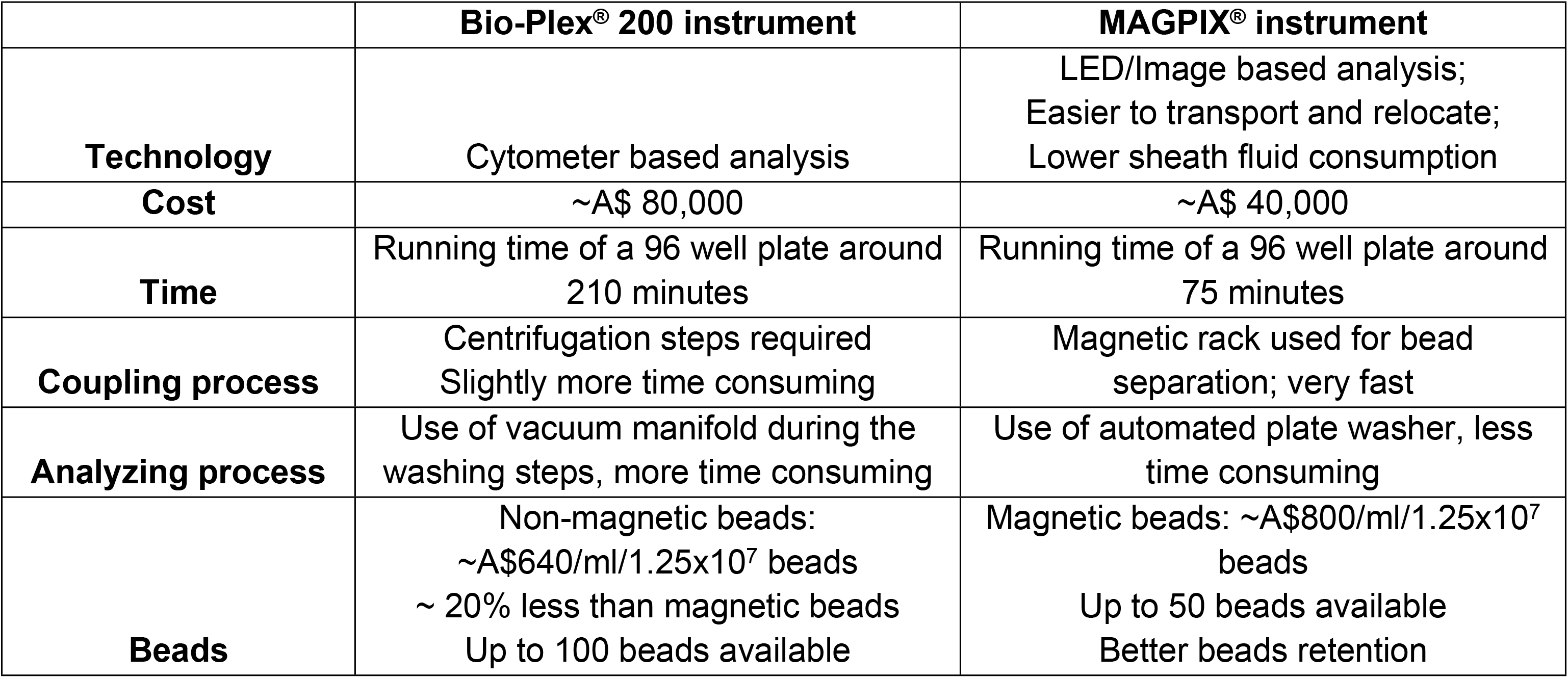
Comparison of the two platforms commonly used for Luminex bead-based assays.

|  | Bio-Plex® 200 instrument | MAGPIX® instrument |
| --- | --- | --- |
| **Technology** | Cytometer based analysis | LED/Image based analysis;<br>Easier to transport and relocate;<br>Lower sheath fluid consumption |
| **Cost** | ~A\$ 80,000 | ~A\$ 40,000 |
| **Time** | Running time of a 96 well plate around<br>210 minutes | Running time of a 96 well plate around<br>75 minutes |
| **Coupling process** | Centrifugation steps required<br>Slightly more time consuming | Magnetic rack used for bead<br>separation; very fast |
| **Analyzing process** | Use of vacuum manifold during the<br>washing steps, more time consuming | Use of automated plate washer, less<br>time consuming |
| **Beads** | Non-magnetic beads:<br>~A\$640/ml/1.25x10<sup>7</sup> beads<br>~ 20% less than magnetic beads<br>Up to 100 beads available | Magnetic beads: ~A\$800/ml/1.25x10<sup>7</sup><br>beads<br>Up to 50 beads available<br>Better beads retention |

For future use and development of the assay, we recommended that a reference laboratory provide both protein-coupled beads and a positive control, along with a Standard Operating Procedure for the assay. All protein-coupled beads should be tested for stability and researchers provided with an expiry date for their use, in addition to checking the performance of the standard curve before each use. This should ensure repeatable and comparable measurements are generated between different research groups. A key focus of *P. vivax* serology efforts should be to develop a standard WHO reference reagent for *P. vivax* that is available to any research group worldwide.

Whilst these results were obtained in the context of *P. vivax*-specific IgG responses in individuals from malaria-endemic areas, the large panel of proteins used and consistent results obtained for all proteins suggest these results can be applied to guide studies in other fields. Luminex xMAp^®^ technology has been used to measure antibody responses against other infectious pathogens, such as HIV and influenza [17, 18], to a variety of vaccine antigens such as tetanus toxoid [19], and more recently to SARS-CoV-2 [6–8].

## Acknowledgements

We wish to acknowledge the extensive field-teams in Thailand, Solomon Islands and PNG that originally collected samples in the studies that were used for this project. We thank Connie Li-Wai-Suen for providing the R code for the standard curve transformation.

## List of Supporting Information Files

**Figure S1**. Comparison of IgG antibody levels against 12 *P. vivax* proteins when run at WEHI compared to CWRU: raw MFI values.

**Figure S2**. Comparison of IgG antibody levels against 12 *P. vivax* proteins when run at WEHI compared to CWRU: converted RAU values.

**Figure S3**. Stability of protein-coupled magnetic beads over 9-months. The original coupled beads were tested at every week for 2 months after coupling, then again at 9 months post-coupling. The MFI of the standard curves are presented (S1 = 1/50, then 2-fold serial dilution). New vials of secondary antibodies were opened on 19/02/19, 26/02/19 and 08/03/19. Protein PVX_094255 (WGCF construct) was not tested in this experiment.

